# Hypotheses and models linking epigenetic transgenerational effects to population dynamics: Exploring oscillations and applications to wildlife cycles

**DOI:** 10.1101/2020.05.05.079129

**Authors:** David Juckett

## Abstract

Epigenetic transgenerational mechanisms underpin the imprinting of gamete origin during reproduction in mammals but are also hypothesized to transmit environmental exposures from parents to progeny in many life forms, which could have important consequences in population dynamics. Transgenerational hypotheses embody epigenetic alterations occurring in gametes, embryonic somatic cells, and embryonic primordial germ cells because most of the epigenome is erased and reconstituted during development. Four scenarios are described in this paper encompassing somatic and germline effects where each of these is either non-propagating or propagating in time. The non-propagating effects could result from environmental impulses such as toxicants, weather, epidemics, forest fires, etc. The propagating effects could result from continuous signals such as climate variations, food web abundances, population densities, predator numbers, etc. Focusing on the propagating mode, a population growth model is constructed incorporating the intrinsic delays associated with somatic or germline effects. Each exhibit oscillatory behavior over a wide range of the parameter space due to the inherent negative feedback of such delays. The somatic (maternal) model oscillates with a period of ∼6 generations while the germline (grandmaternal) model oscillates with a period of ∼10 generations. These models can be entrained by oscillatory external signals providing that the signals contain harmonic components near the intrinsic oscillations of the models. The 10-generation oscillation of the germline-effects model is similar to many wildlife cycles in mammals, bird, and insects. The possibility that such a transgenerational mechanism is a component of these wildlife cycles is discussed.

## 1 Introduction

**T**Here is a growing interest in the concept of environmental exposure effects being transmitted across generations (e.g., Bonduriansky et al. (2012); Skinner (2015); Walsh et al. (2016); Safi-Stibler & Gabory (2020)). Effects that may be transient and wash out in a few generations or effects that may track with environmental variations are most likely to involve the epigenome because genetic mutations don’t have such plasticity. Such effects can be referred to as epigenetic transgenerational signaling. If the epigenetic alterations remain stable over several unexposed generations, then this is often referred to as epigenetic transgenerational inheritance (Jablonka & Raz (2009); Skinner et al. (2012); Sharma (2017)). The epigenetic modifications that must be renewed each generation have profound implications for population dynamics because they provide mechanisms for maintaining species niches under the pressures of varying external forces. These types of epigenetic modifications are the focus of the models presented here.

Direct evidence for epigenetic changes across generations has been demonstrated in several organ systems (e.g., Buiting et al. (2003) in neural disorders; Anway et al. (2005) in fertility changes from endocrine disruptors; Alm et al. (2017) in muscle protein changes; and the Millership et al. (2019) review of effects on metabolism). Inferred epigenetic effects have been demonstrated by Plaistow et al. (2006) in intergenerational effects observed after food restrictions in soil mites.

Mechanisms for epigenetic programming have been reviewed by many (e.g., Jablonka & Raz (2009); Gilbert & Epel (2009); MacDonald (2012); Skinner (2016)). Epigenetic reprogramming has been shown to occur in the primordial germ cells of mouse embryos and while the purposes for some epigenetic changes are known, many are not (Matsui & Mochizuki (2014); Kurimoto & Saitou (2019)). In addition to germline reprogramming, the somatic epigenome of the zygote is mostly erased and is reprogrammed in the somatic cells of the embryo after implantation (Zeng & Chen (2019); Safi-Stibler & Gabory (2020)). This offers an additional mechanism for intergenerational transmission. In the germline case, an epigenetic alteration in the F0 generation would be manifested in the F2 generation. In the case of a somatic cell alteration, the effect would appear in the F1 generation.

In general, the transmission of effects across generations could be explained by two separate mechanisms. The first would be a passive process where environmental molecules or energy fluxes directly interact with male or female gametes of the F0 adults prior to zygote formation or directly interact with the germline in the early embryo (Skinner, 2016). The second would be an active process where an environmental signal is detected by the mother and transmitted to either the embryonic somatic cells (maternal effect) or to the germline (grandmaternal effect). When either somatic or germline epigenetic effects occur throughout a population then it is reasonable to assume that they would influence population growth dynamics due to widespread variations in fecundity, fertility, and general vigor. The active transmission process allows species to modify growth dynamics to maintain their niche.

Maternal effects on population growth dynamics were addressed by Ginzburg & Taneyhill (1994) and reviewed by Inchausti & Ginzburg (2009) and Ginzburg & Krebs (2015) as a mechanism capable of driving cycles in various wildlife species. Maternal effects have also been invoked by others in wildlife cycling (e.g., Rossiter (1994); Benton et al. (2001); Rakyan et al. (2003); Plaistow et al. (2006); Anderson & Gillooly (2017); Boonstra et al. (1998); Krebs et al. (2018) and references therein). The underlying mechanisms, however, have not been definitively demonstrated. The very nature of maternal influences during reproduction highlights the importance of generation interval in population dynamics rather than simple time units. This has been referred to as the 4th dimension in biology (Ginzburg & Colyvan (2004); Ginzburg & Damuth (2008)), offering a more consistent metric for understanding population dynamics.

If the maternal signaling occurs within the germline rather than the somatic cells of the embryo, then the effects would skip the F1 generation and affect the F2 generation. This can be referred to as a grandmaternal effect. Transgenerational inheritance effects through the germline have been widely explored (see reviews of Jablonka & Raz (2009); Skinner (2016)). The possible roles of this effect in health, disease, and aging have also been explored (Sharma (2017); Nilsson et al. (2018); Safi-Stibler & Gabory (2020)). These studies have focused on semi-permanent germline modifications that can be long-lived. Epigenetic germline effects that must be renewed every generation have received little attention. Indirect evidence for renewable germline epigenetic reprogramming has been presented for grandmaternal inheritance in human cancer predisposition (Juckett & Rosenberg (1993); Juckett & Rosenberg (1997); Juckett (2007); Juckett (2009)). This grandmaternal mechanism could also influence population growth dynamics but it, too, has not received much attention. Sinclair et al. (2003) provided evidence for longevity and senescence variations in the snowshoe hare cycle and suggested that transgenerational effects, including grandmaternal effects could be at play.

In this paper, maternal and grandmaternal effects are explored conceptually and in mathematical models. In the next section, the foundational postulates are given, followed by hypotheses for this work, and then the conceptual model that outlines propagating and non-propagating modes. A mathematical population model that can embody either the maternal or grandmaternal propagating modes is presented in the subsequent section. In the Results section, the intrinsic oscillations of the mathematical model are explored including the ability to be entrained by external oscillations. With a focus on the 10-generational cycle of the germline grandmaternal model, comparisons and predictions are made to various wildlife cycles. A heuristic example is also provided showing the similarity between the snowshoe hare and ruffed grouse cycles and a possible entraining signal in the form of the 10-yr component of Pacific Decadal Oscillation. Finally, the model and results are discussed in the context of populations cycles in nature.

## 2 Postulates, Hypotheses, and Conceptual Model

### 2.1 Postulates

To form a basis for hypothesis generation and model building, the postulates that form the underlying assumptions of this work are as follows:

1. Populations of organisms must respond to changing environments to prevent overpopulation when resources are limited and to prevent underpopulation and loss of niche when resources are plentiful.
2. To facilitate species survival, responses to environmental cues must be coordinated among individuals in the population.

The first postulate was probably expressed initially in its general form by Chitty (1957) who proposed that species can self-regulate their populations in response to the environment before population numbers reach the limit of available resources. The second postulate is a natural extension of the first postulate and is assumed to be a product of evolution. It would allow population members to act in synchrony to adapt to the environmental pressures for the betterment of the species and not just the individual. The second postulate also allows an exploration of transgenerational effects through the study of population dynamics.

### 2.2 Hypotheses

A cascade of hypotheses, built upon the postulates, is presented to breakdown the steps for maternal and grandmaternal processes modulating population numbers.

1. The response to changing environments would be expressed as changes in progeny fertility and/or vigor.
2. Transgenerational epigenetic reprogramming provides the mechanism to respond to changing environments.
3. Epigenetic reprogramming occurs in the embryo, driven by signals received from the mother.
4. Embryonic epigenetic reprogramming can occur in either the somatic embryo or the embryonic primordial germ cells.
5. Intra-species communication occurs between fecund females via mechanisms such as pheromone exchange to coordinate environmental responses.

The last hypothesis, as well as the corresponding postulate, are presented as justification to study transgenerational effects in the context of population dynamics. Neither are explored further in this paper because an in-depth look at mechanisms and data would be required.

### 2.3 Conceptual Model

Four transgenerational epigenetic mechanisms are shown in Table 1. The targets listed in the table are the genomes during the time they are being reprogrammed with epigenetic markers after implantation of the zygotes. The course of events are shown in the cartoons of Figures 1A & 1B. The environmental signals are depicted by solid black arrows. For the continuous modes, only two of these arrows are shown allowing clarity in how effects propagate through the generations and how they wash out. The continuous environmental signals would be present at each generation and it is assumed that epigenetic reprogramming must be renewed each generation and would be dictated by the instantaneous environmental signal present at that time.

**Table 1:** Overview of epigenetic transgenerational modes for populations.

| Epigenetic mode | Stimulus | Target |
| --- | --- | --- |
| Somatic non-propagating | Pulse | Somatic embryo |
| Somatic propagating | Continuous | Somatic embryo |
| Germline non-propagating | Pulse | Primordial germ cells |
| Germline propagating | Continuous | Primordial germ cells |

**Figure 1a.**
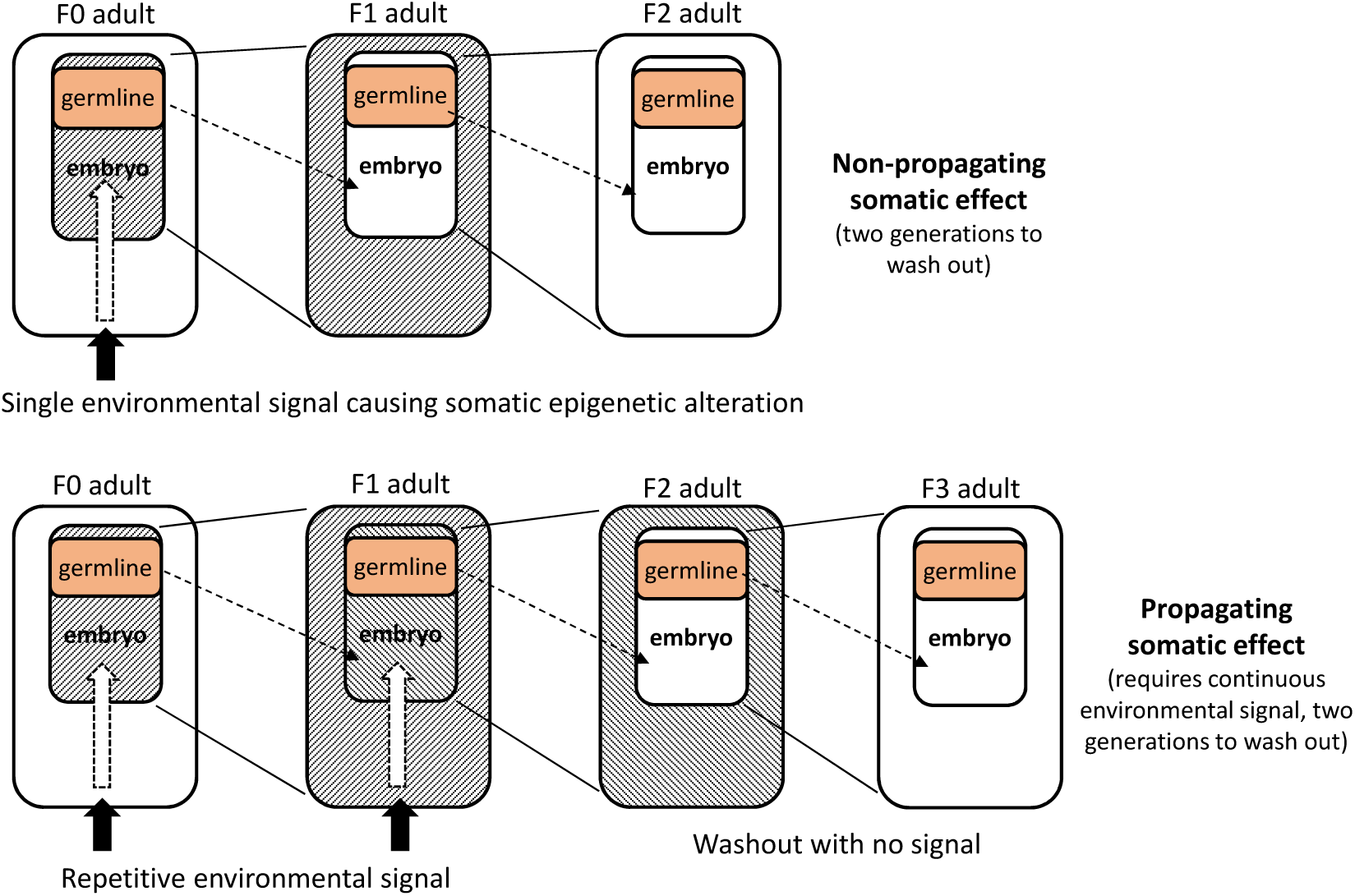
Conceptual representation of non-propagating and propagating somatic epigentic transgenerational effects.

**Figure 1b.**
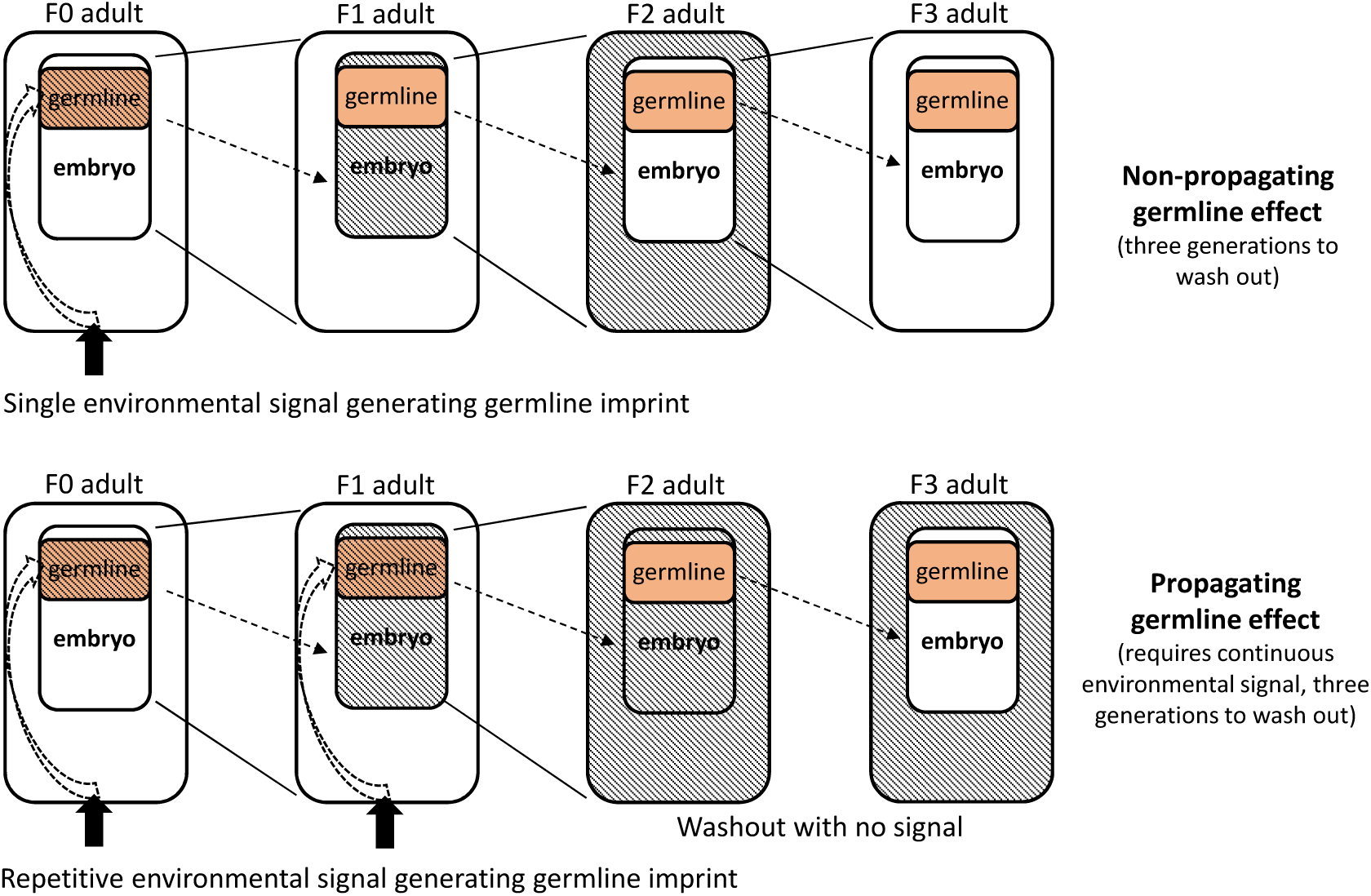
Conceptual representation of non-propagating and propagating germline epigentic transgenerational effects.

The somatic (maternal) effect is depicted in Figure 1a. The environmental signal is detected by the F0 female who, in turn, would impart an effect on the somatic cells within the growing embryo during epigenome reprogramming. This epigenetic modification is expressed within the F1 adult, as shown by the hatched area. Since the germline is unaffected, the embryo within the F1 adult does not receive the specific environmental-derived epigenetic signal. In the absence of additional environmental signaling (non-propagating case), the F2 adult epigenome returns to F0 adult status. In the presence of continuous environmental signaling (propagating case), the somatic embryo in the F1 adult also receives the signal and the F2 adult is affected. This would continue as long as the signal is present but would wash out one generation after the affected generation, as shown in Figure 1A. The inherent time delay for population modeling is one generation for a signal to take effect and another generation to wash out.

The germline (grandmaternal) effect is depicted in Figure 1b. The environmental signal is detected by the F0 female who, in turn, would impart an effect on the germline cells within the growing embryo during epigenome reprogramming. This modification is not expressed within the F1 adult but is carried by the gametes of the F1 generation. The F2 adult expresses the modification and for the single impulse, non-propagating case, washout occurs in the F3 generation. In the presence of continuous environmental signaling (propagating case), the germline cells are affected in each generation. The epigenetic effect would always lag behind the environmental signal by two generations. The inherent time delay for population modeling in the germline effect scenario is two generations for the signal to take effect and one more generation to wash out. In practice, the propagating mode would be the response to varying environmental signals that may be oscillatory or chaotic in nature and therefore would not wash out, per se. Nevertheless, the delay between changes in the signal and changes in the population would be one generation for the somatic (maternal) mechanism and two generations for the germline (grandmaternal) mechanism. These are delays that can be easily introduced into population growth models.

To provide context for these propagation modes, the non-propagating transgenerational effects may occur with sporadic or non-continuous processes, such as exposures to toxicants, disease epidemics, forest fires, catastrophic weather events, etc., and would tend to be highly localized and relatively short-lived in time. The non-propagating transgenerational process could serve to briefly modulate reproduction and vigor to help the local population maintain its niche over the short term.

The propagating transgenerational effects are of greater interest to population biology. These are likely to be induced by continuous processes, such as predation intensity (including parasites), food web abundance, nesting availability, population density, climate variations, weather in its various forms, geomagnetic variations, and extraterrestrial fluxes such as ultraviolet radiation and cosmic rays. Recent work documents that just the fear of predation can also affect population reproduction and survival (Macleod et al., 2018). Some of these are local and some are more global in nature. The propagating transgenerational process could serve to modulate reproduction and vigor to help broad populations maintain niche and viability over the long term.

## 3 Mathematical Population Model

Cycles in various wildlife species have been explored for many decades and many mathematical models have been constructed. Both differential equations and discrete difference equations have been used for single-species or multi-species predator-prey models. To provide negative feedback, incorporation of a time delay was introduced in difference equations by Hutchinson (1948), in differential equations by Cunningham (1954), and was explored more by May (1972) and many others. Negative feedback readily leads to robust limit cycles spanning a wide range of parameter values. While the stability of time-delay models has been repeatedly shown, the source of such delays has not always been obvious. The transgenerational inheritance hypothesis may provide a source for many of these delays.

To explore the epigenetic transgenerational hypothesis in its simplest form, a two-parameter single species population model was numerically solved for various parameter values. This model uses a logistic-type term to provide density dependency, which accounts for resource limitations and predation effects by incorporating a set point for stable population numbers. The species would attempt to regulate its population numbers around this set point. The model can be formulated with either a one-generation delay or a two-generation delay. This provides an intrinsic negative feedback, which has the side effect that it naturally leads to robust population cycles for certain regions of the parameter space. The model is also easily expanded to allow the introduction of a synchronizing external oscillation, such as regional climate variations.

### 3.1 Single-species, two-parameter forumulation

In the spirit of not over-parameterizing the problem (Ginzburg & Jensen, 2004), a single-population formulation was chosen with a minimum of parameters. The growth of a single population can be represented by the following differential equation:

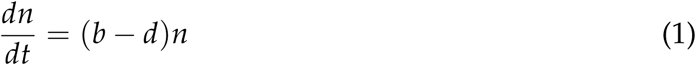

For discrete times units of generations, this can be rendered as a difference equation:

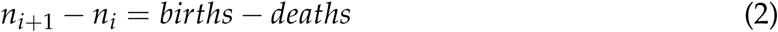

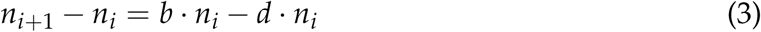

where *n* is the population count, *b* and *d* are the birth and death rates, and *i* represents the generation count. In a steady-state scenario, birth and death rates would be equal and the population stable. The difference between the birth and death rates can be replaced by two alternate parameters; the first being the net reproduction rate, *a*, under ideal conditions and the second being a perturbation factor, Δ, where Δ would be zero in the steady state and would vary around zero to yield both population increases and decreases. Together, the factor (*a ·* Δ) yields an effective growth rate. This is given by:

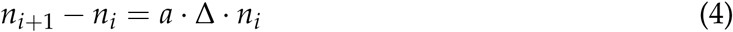

Eq. (4) can be considered a canonical representation of population dynamics because of its inherent simplicity.

To provide stability to the population dynamics under this definition of *a*, the value of Δ needs to be no greater than 1. One possible choice for Δ would be:

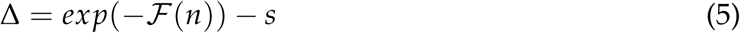

where the exponential provides a term bounded by 0 and 1 for *ℱ* (*n*) ≥ 0. Using the exponential function allows rapid recovery when *ℱ* (*n*) is low and a slow change when *ℱ* (*n*) is high. This allows robust behavior over wide-ranging values of *n*. (The exponential can be replaced by other logistic style terms, see the Discussion for examples.) The value of *s*, defined for 0 < *s* ≤ 1, provides the threshold between population gain and loss, which is a set point determined by the long-term (many generation) balance of factors such as; prey vigor, prey fertility, resource availability, and predation. In general terms, the value of *s* can be considered a normalized environmental stress factor averaged over time.

To incorporate the maternal or grandmaternal effect in the simplest manner, *ℱ* (*n*) is made equal to the population one or two generations earlier. Eq. (5) then becomes:

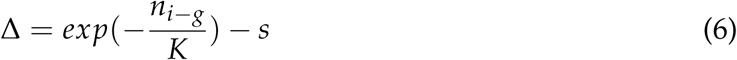

where *g* has a value of one for the maternal effect and a value of two for the grandmaternal effect and *K* is the mean population size over a span of many cycles. Combining Eqs. (4 & 6), yields the final form of the two-parameter equation that represents one possible population growth model containing density dependence and a set point.

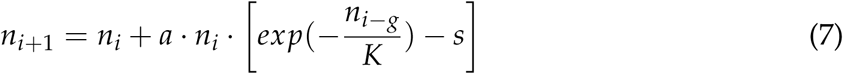

This equation falls into the large class of delayed density dependent models with the advantage that the equilibrium point, *s*, is confined to the easily explorable range 0 < *s* < 1.

It is important to make the connection between Eq. (7) and Figure 1. In Eq. (7), the size of the (*i* + 1) generation is the result of the (*i* − *g*) generation acting on the *i* generation. In the case of *g* = 2, the F2 adults exhibit altered characteristics inherited from the F0 generation which don’t cause a populations change until the F3 adults. This is the difference between effects on individuals and effects on populations. This is consistent with observations on reproduction changes and timing in snowshoe hare populations across their 10-yr cycle (Cary & Keith, 1979).

When Δ is zero the population is at steady state. Setting Δ = 0 in Eq. (6), rearranging and taking the logarithm yields:

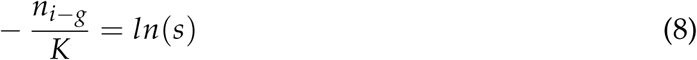

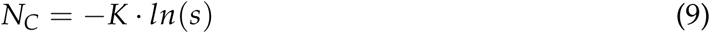

The value *n* at steady state, denoted *N*_*C*_, is the size of the population with balanced births and deaths and could be interpreted as the size of the population needed for equilibrium when the mean environmental stress is *s*. This could also be referred to as the mean carrying capacity, which varies with environmental conditions. It is clear from Eq. (9) that the value of *ln*(*s*) is always negative and as *s* increases *ln*(*s*) approaches zero consistent with a reduced carrying capacity.

### 3.2 External synchronization

The model has intrinsic negative feedback and, therefore, exhibits natural oscillations for certain ranges of the two parameters, *a* and *s*. Such model oscillators typically can be modulated or entrained by external oscillations (Juckett, 2010). This can be introduced into Eq. (7) by modulating the environmental stress parameter, *s*:

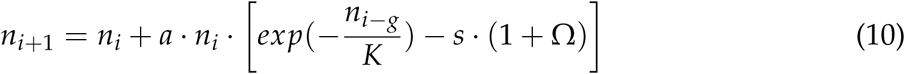

where Ω is an oscillatory driving force represented by a sinusoidal function. In this case, positive Ω effectively increases *s* representing a more inhospitable environment. The three-parameter form of Ω is given by:

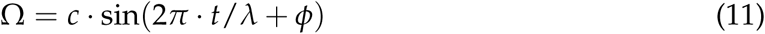

for 0 ≤ *c* ≤ 1, where *t* is time and *c, λ, ϕ* represent the three degrees of freedom; amplitude, period, and phase. By using (1 + Ω) in Eq. 10., it provides a modulating function that oscillates around a value of 1.0. Equation 11 can also be represented by a superposition of harmonic waveforms, which is typical of most environmental signals of temperature, pressure, humidity, sunny days, etc.

Under the assumption that the environmental signal, embodied by Ω, acts in the (*i* − *g*) generation to modulate the species response to the changing external conditions, Eq. 10 takes the following form:

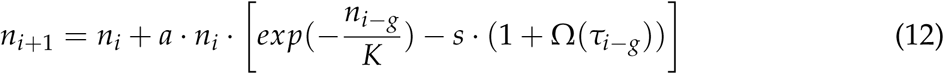

where, *τ*, is a variable restricted to discrete increments representing the mean time for one generation, *i*, and the parameters *c* and *φ* are suppressed for clarity.

Since formulations with negative feedback, like Eq. 7, exhibit various behaviors, the introduction of an external oscillation as shown in Eq. 12 can have various affects. When the model has parameter values that already lead to oscillations, the presence of an external signal can entrain the native oscillation into a constant phase relationship to the external signal. In regions where the model exhibits steady state dynamics, the external signal can pump the model and cause cycling. These effects depend on the amplitude of the external oscillation and how close the frequency of the external oscillation is to the native resonance of the model.

## 4 Methods

### 4.1 Data

A compilation of data on various species that exhibit population cycling was obtained from an appendix document accompanying the Anderson & Gillooly (2017) publication. Their table included approximate cycling periods (sometimes rounded to integers), generation intervals, method and significance estimate for cycle determination, and source references. Cycles considered statistically significant by the original authors were extracted from the appendix, excluding protists and zooplankton.

Data of the snowshoe hare cycles were obtained from several sources. These are detailed in Appendix A. A compilation series was constructed by scaling each separate series by its root mean square (RMS) amplitude, which allowed comparison across various types of observational units. While such scaling precludes extracting meaningful amplitude variations, it allows a suitable series for detecting the primary frequency component.

Grouse data from Minnesota was obtained from the 2019 Minnesota Spring Grouse Surveys report (Roy, 2019) for the ruffed grouse, spanning 1949 to 2019. Values were digitized from the figure 3 in that report.

**Figure 2.**
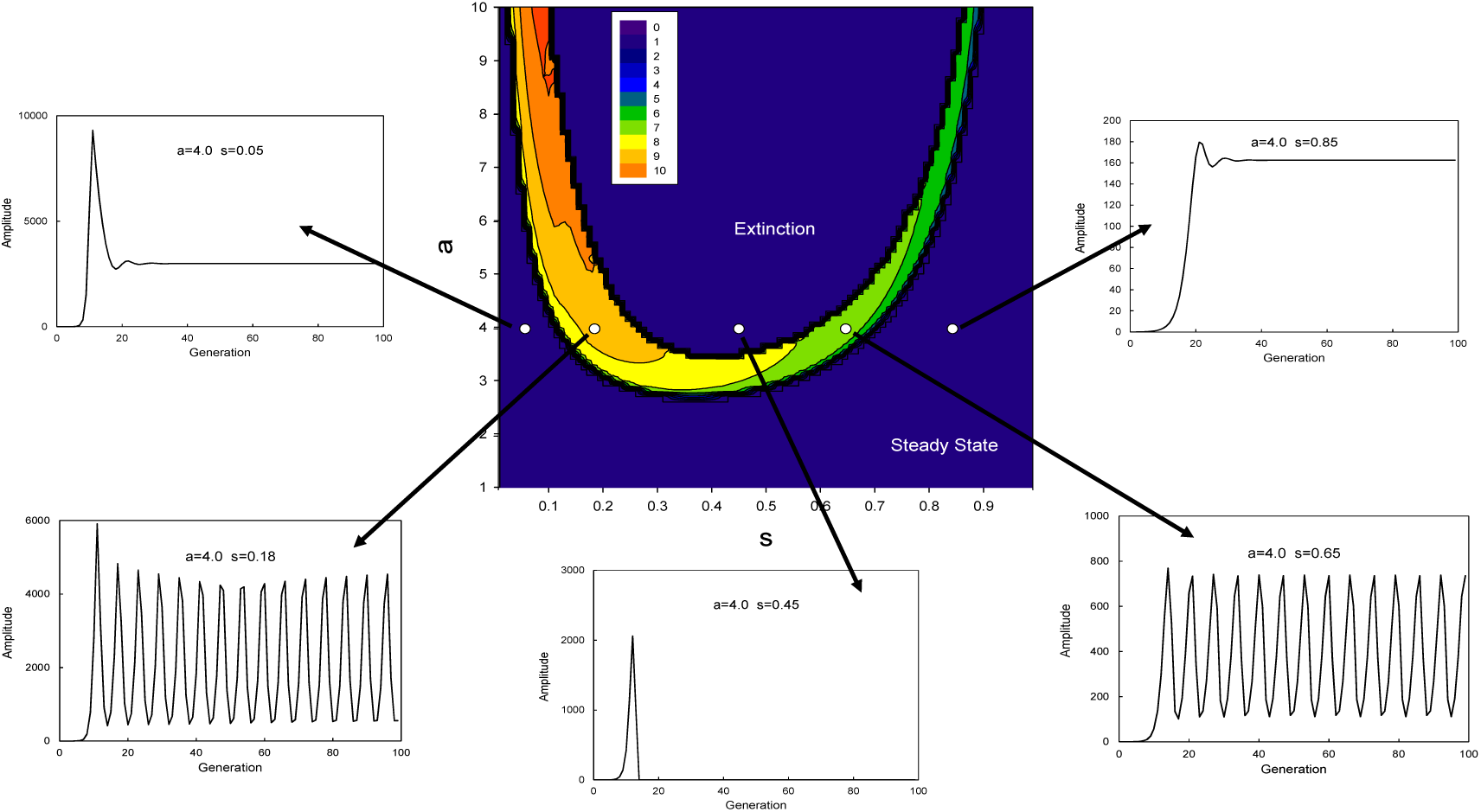
Maternal model (*g* = 1). Amplitude profile of stable oscillations through the region of parameter space defined by 1.0 ≤ *a* ≤ 10.0 and 0.0 < *s* < 1.0. The contour plot was built from evaluating the model dynamics for values of *a* spaced at 0.1 intervals and for values of *s* spaced at 0.01 intervals. Legend indicates *ln*(*amp*) values by color. Amplitude is defined as the difference between maxima and minima values over several cycles at high generation number (i.e., mean peak to trough difference). For steady state and extinction regions, this difference is zero. To accommodate this for the contour plot, *ln*(*amp* = 0) is set equal to zero. Callout figures give examples of model behavior along five regions of the parameter space denoted by open circles. The values of *a* and *s* are given in the callout figures. *K* = 1000.

**Figure 3.**
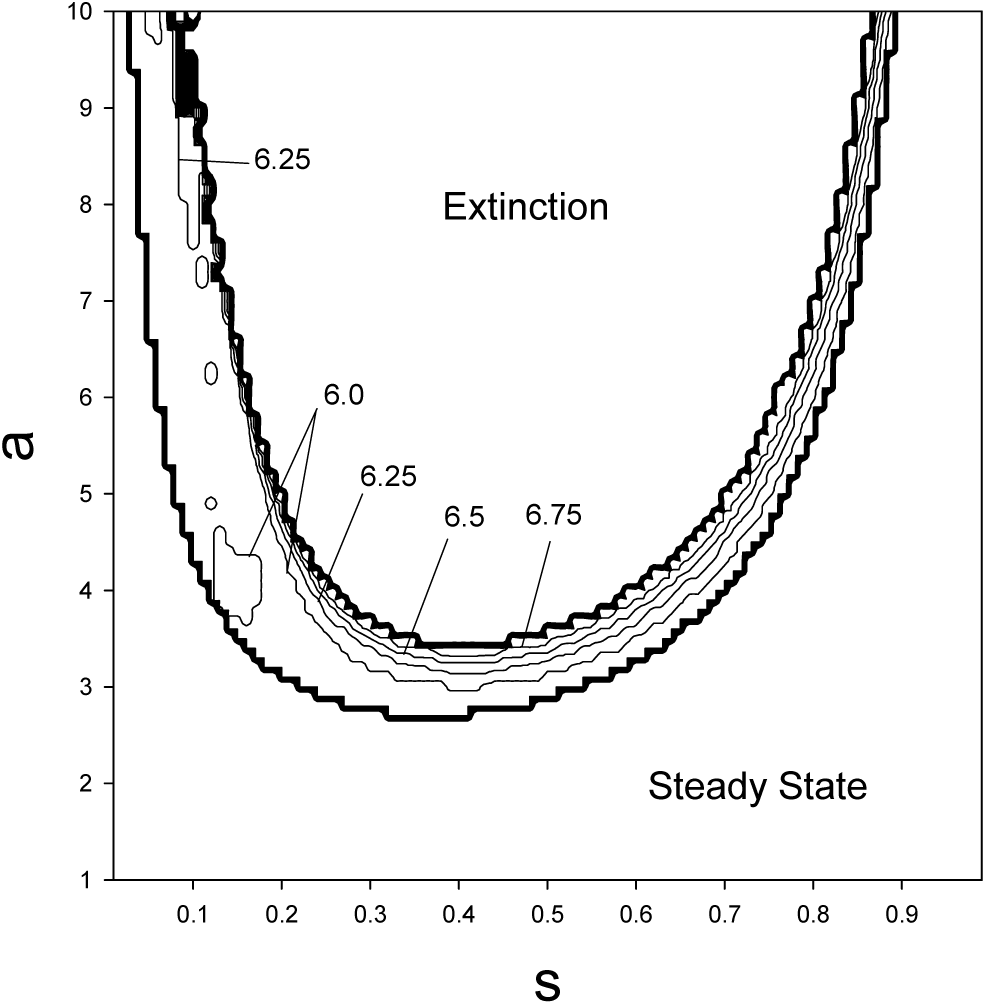
Profile of cycle periods (in generations) within the stable oscillatory region of the parameter space. Contour lines are provided at 0.25 intervals. Oscillation periods vary smoothly across the stable region.

The pacific decadal oscillation (PDO) was obtained from NOAA, downloaded from https://www.ncdc.noaa.gov/teleconnections/pdo/ on January 13, 2020.

### 4.2 Analysis

Evaluations of the difference equation with and without sinusoidal entrainment was performed in the MathCad™ platform. Spectral analysis of time series generally consisted of Fourier analysis and spectral band isolation. Fourier transform analysis was typically performed with zero padding to 4096 points to allow for accurate determination of the band peak locations. In some cases, the Lomb Periodogram was used. Plots were generated in MS Excel™ and SigmaPlot™.

## 5 Results

### 5.1 Model dynamics

The behavior of Eq. (7) was examined over values of *a* from 1.0 – 10.0 and *s* from 0.01 – 0.99 with either no forcing or with single-frequency forcing. Since *a* can be considered the maximal effective progeny size per each member of the population (i.e., number of progeny reaching independent status under optimal conditions), an upper value of ten is deemed sufficient to accommodate most wildlife species, since this would mean a viable progeny size of 20 for a mating pair. The model is stable for much higher values of *a*, which might occur in insects, but for simplicity this was not examined in detail. For each value of *a* and *s* the time series were run out to *i* = 4000 and the region from *i* = 150 − 1000 was evaluated for stable oscillations. The region from *i* = 700 − 1000 was used to extract maximum and minimum values for the differences used to construct Figs. 2 & 4. For each value of *a* and *s*, the sequential evaluation of Eq. (7) was arbitrarily begun with *n*_0_ = 0.1 and the next seven *n* values given by *n*_*i*+1_ = *n*_*i*_ *·* 1.1. These values were away from equilibrium and therefore represented a perturbation leading to compensating population changes.

**Figure 4.**
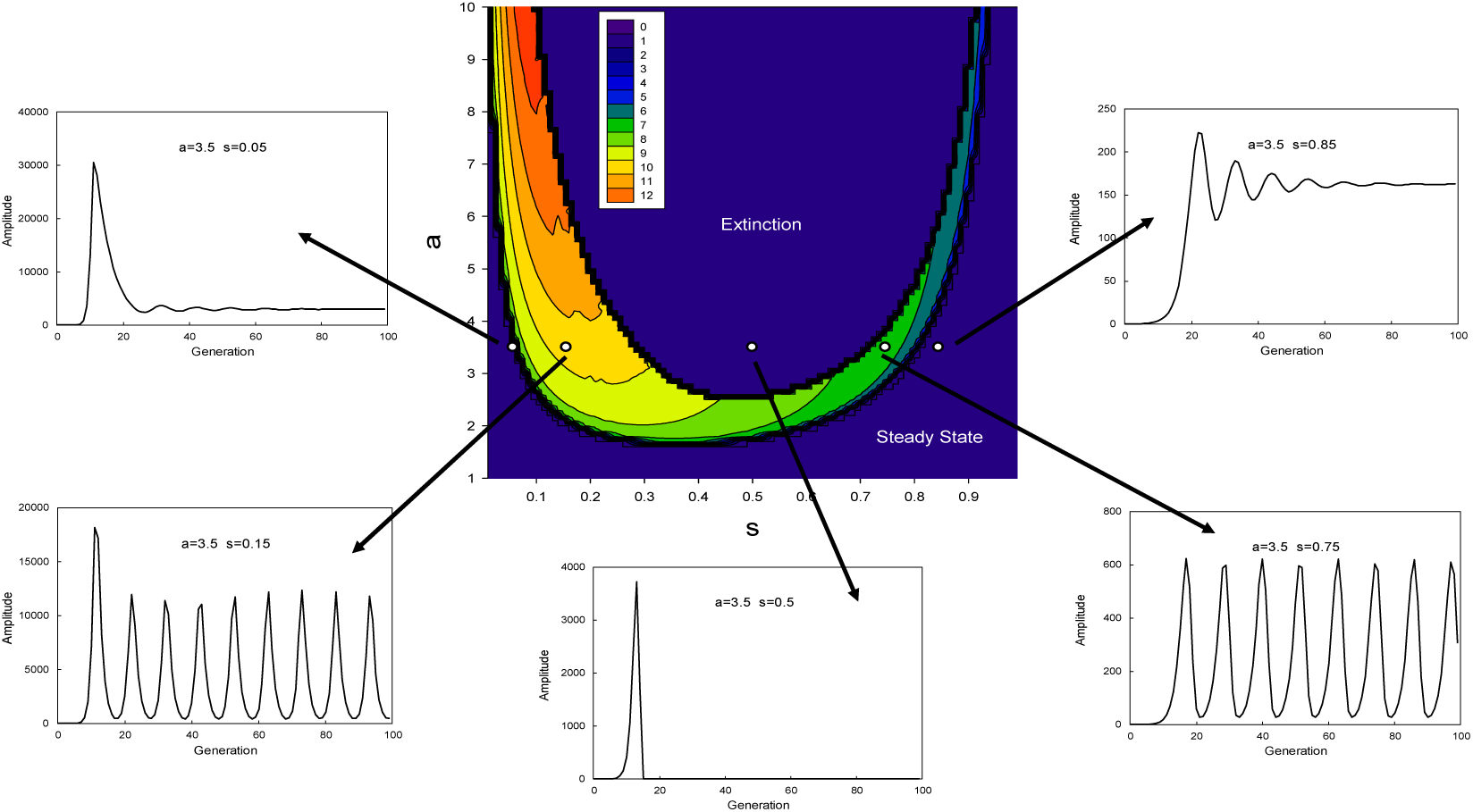
Grandmaternal model (*g* = 2). Amplitude profile of stable oscillations through the region of parameter space defined by 1.0 ≤ *a* ≤ 10.0 and 0.0 < *s* < 1.0. The contour plot was built from evaluating the model dynamics for values of *a* spaced at 0.1 intervals and for values of *s* spaced at 0.01 intervals. Legend indicates *ln*(*amp*) values by color. Amplitude is defined as the difference between maxima and minima values over several cycles at high generation number (i.e., mean peak to trough difference). For steady state and extinction regions, this difference is zero. To accommodate this for the contour plot, *ln*(*amp* = 0) is set equal to zero. Callout figures give examples of model behavior along five regions of the parameter space denoted by open circles. The values of *a* and *s* are given in the callout figures. *K* = 1000.

### 5.2 Native oscillations - maternal and grandmaternal effects

The region of spontaneous, stable oscillations within the [*a, s*] parameter space is shown in Figs. 2 & 3 for the one-generation delay (*g* = 1) model and in Figs. 4 & 5 for the two-generation delay (*g* = 2) model. Depending on the parameter values, different behaviour ensued as shown in the Figs. 2 & 4 where the natural logarithm of the cycle peak to trough differences is shown as a contour plot. The differences can exceed 10^4^ − 10^6^ in some regions, which is well beyond the range seen in the natural environment, with the possible exception of insect outbreaks. The U-shaped domain of stability exhibits a uniform, but rapid increase to high peak trough differences in the region of low *s*. In the lower part of the *a* range, the populations exhibit steady state levels of no growth and no oscillation. Above the ‘U’ region of stable oscillations, populations rapidly go to extinction. In Figs. 2 and 4, five examples are shown from different regions of the parameter space showing the initial transitions to stability, oscillation, or extinction.

Fig. 3 is the contour plot of the Fourier spectra for each choice of *a* and *s* over the region from i = 500-4000 in each series in the one-generation delay model. The typical period is approximately 6 – 6.75-generation intervals and is independent of cycle amplitude, birth rate, and set point. These cycle periods are consistent with the values predicted and demonstrated by Ginzburg & Colyvan (2004) using a different mathematical approach. Fig. 5 is the contour plot for the two-generation delay model calculated in the same manner. The typical period is between 10 – 11.5 generations over the broad U-shaped region of stable oscillation. The periods for these models are within expectation for time-delay models (May, 1973), but in this case, periods are in generations, not time.

**Figure 5.**
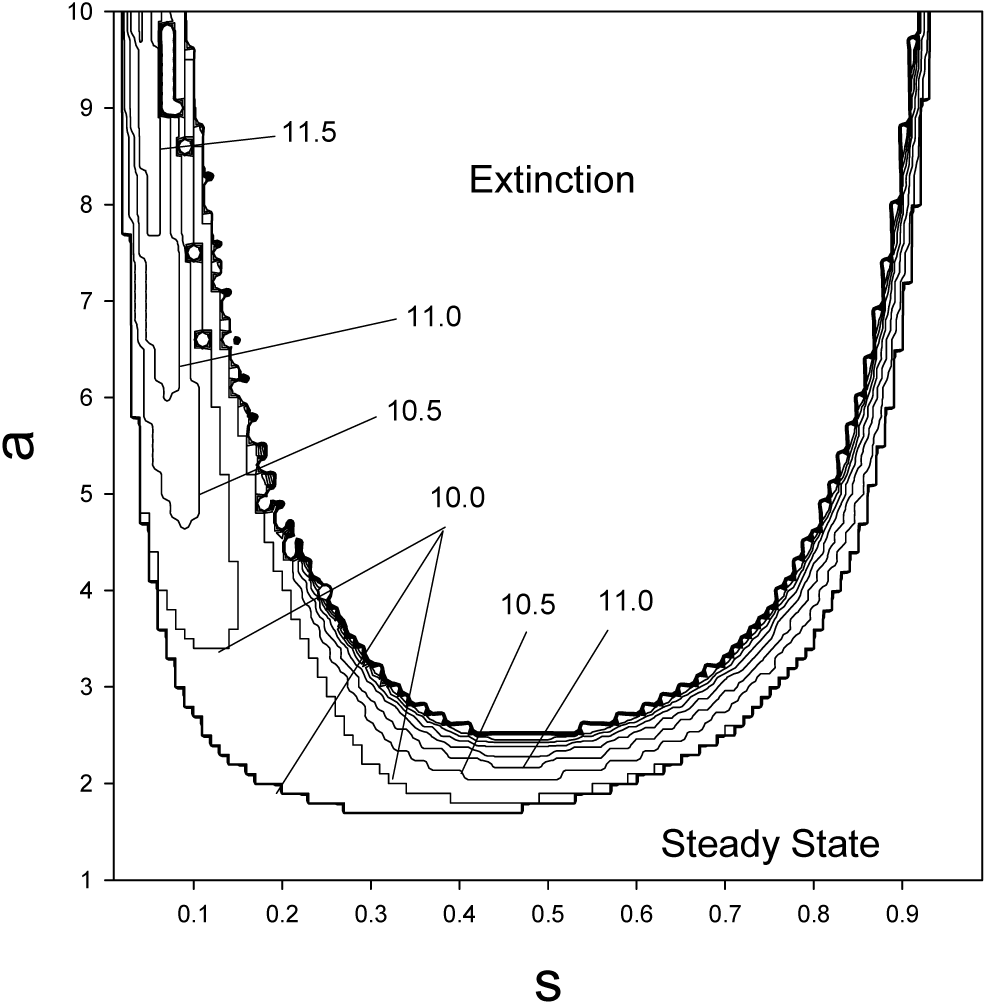
Profile of cycle periods (in generations) for the grandmaternal model within the stable oscillatory region of the parameter space. Contour lines are provided at 0.5 intervals. Oscillation periods vary smoothly across the stable region.

If the maternal and grandmaternal effects are both operating in a particular species then the model can be adapted by changing the numerator of the exponential in Eq. 7 to (*n*_*i*−2_ + *n*_*i*−1_). In this case, the model has an intrinsic oscillation with cycle periods near 8.0 generations and the amplitudes and frequency responses over the parameter space resemble those of Figs. 2 to 5. If there are varying degrees of the two effects, then the intrinsic cycle lengths can be anywhere between 6.0 and 10 generations.

The area labeled as ‘Extinction’ in Figs. 2-5 may be an artifact of the simplicity of the mathematical model in which overcompensation after the first peak drives the population to zero. While different definitions of delta do not eliminate this region, mechanistic models might modulate the rapid population decrease in such a way that extinction is avoided, but that is beyond the scope of this paper.

In summary, results of Figs. 4 and 5 for the grandmaternal model indicate that the fundamental cycle period for the model can be approximated by:

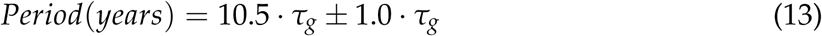

where *τ*_*g*_ represents the mean inter-generational interval for the species in question. To predict the cycle period in years, one only needs to know the mean value of *τ*_*g*_. A similar equation can be constructed for the maternal model.

### 5.3 Synchronized oscillations - example

Mathematical models with stable limit cycles over a broad range of parameter values, like Eq. (7), can be entrained by an external oscillation provided the amplitude is sufficient and the frequency is close to the natural frequency of the model (Glass and Mackey, 1979).

Entrainment is a common occurrence in nature for external signals like the day/night cycle, the seasonal cycles, or the phases of the moon.

In the two-generation delay model (g=2), entrainment is examined for the [*a, s*] value of [3.5, 0.2] using Eq. (11), which is the reformulation of Eq.(7) with the incorporation of a sinusoidal modulation of the *s* parameter. A period of 9.9 yr was chosen for *λ* within the external signal Ω (Eq. (11)). This is close to the intrinsic cycle period of the model under the assumption of one generation per year. The amplitude of the external oscillation required for phase-locking is explored in the example given in Fig. 6. At low Ω amplitude (c=0.00005), the model oscillates with the initial arbitrary phase of approximately −0.3 radians, regardless of the phase of the driver, Ω. As the amplitude of Ω increases, the phase profile of the model becomes a function of the phase of Ω until at sufficient amplitude the model is locked onto the driving sinusoid, often with some constant phase delay. It is noteworthy that the amplitude at the point of phase lock is quite small, indicating a fluctuation as low as approximately 3.0 % can entrain the system.

**Figure 6.**
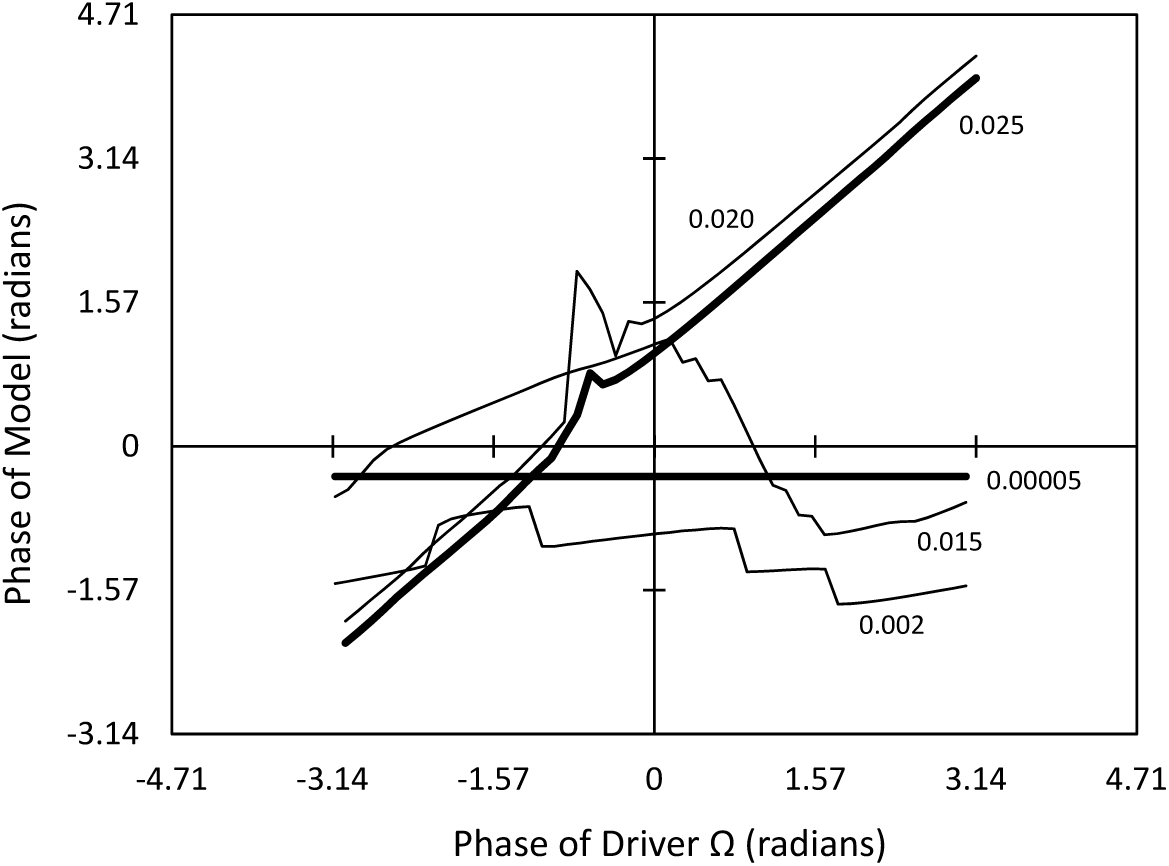
Amplitudes required for phase-locking of the externally driven model (Eq. (12)) to sinusoidal signals of varying amplitudes. This example used [*a, s*] values of [3.5, 0.2] *a* = 3.5, *s* = 0.2, *g* = 2, and *λ* = 9.9 yr. Values next to each line are the amplitudes of the sinusoidal function, Ω. The constant phase just below zero radians is the arbitrary phase of the un-entrained model. Nearly full entrainment is shown in the diagonal bold line with amplitude 0.025.

Using the same values of *a* and *s* as in Fig. 6, the amplitude of Ω that leads to full entrainment was determined for cycle periods surrounding the native cycle period of 10 generations. This is shown in Fig. 7a where amplitude is plotted versus 1/*λ* or frequency. The amplitudes required for entrainment increase with distance from the native frequency of the model, Eq. (7). Taking the inverse of amplitudes generates the equivalent of a bandpass filter with a high Q-value (Fig. 7b) that can be used to determine the influence of external, environmental signals. This bandpass filter is only shown for the [*a, s*] values of [3.5, 0.2], but this basic shape exists throughout the region of stable cycles.

**Figure 7.**
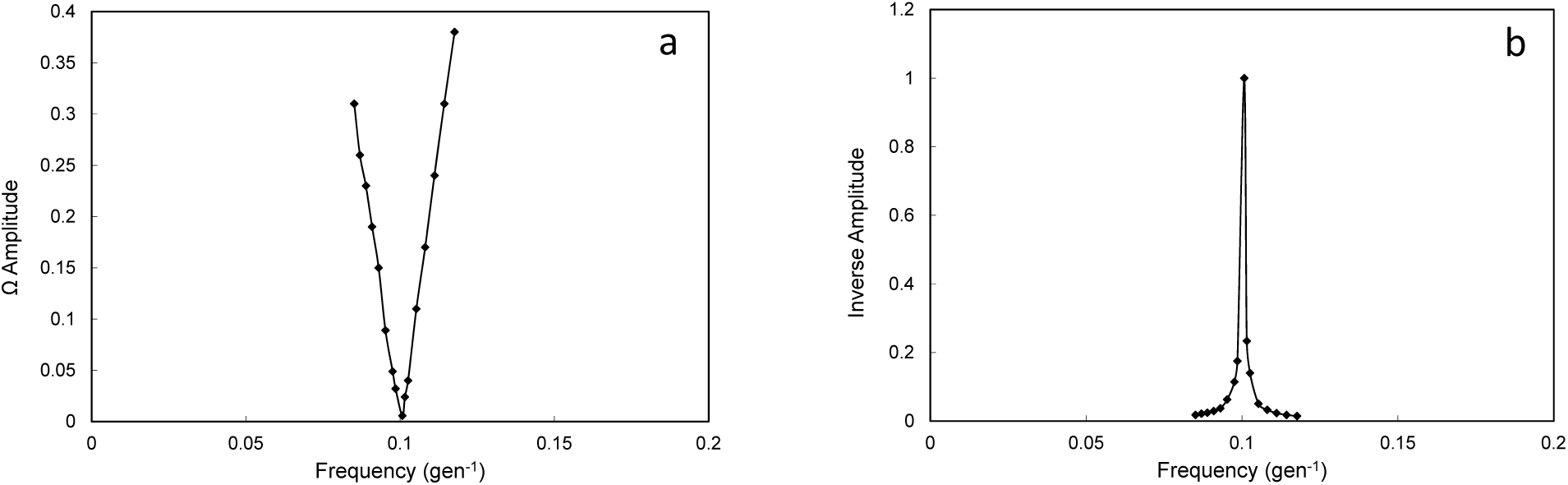
Plot of amplitude **(a)** and inverse amplitude normalized to a maximum value of 1.0 **(b)** necessary for full entrainment of the model at the frequencies shown for external sinusoidal oscillations. The values of parameters [*a, s*] were [3.5, 0.2].

Using the Ω amplitude of c=0.05 and the Ω lambda of 9.9 generations (yr), the contour plots of ln(amp) and cycle periods are shown in Figures 8 and 9, respectively. This forcing oscillation can cause stable cycles in most portions of the parameter space that were shown to contain steady state dynamics in Fig. 5. Thus, this shows both pumping and entrainment. Over the high *a* and low *s* region the cycle periods have remained mostly unchanged, requiring higher amplitude Ω to be entrained.

**Figure 8.**
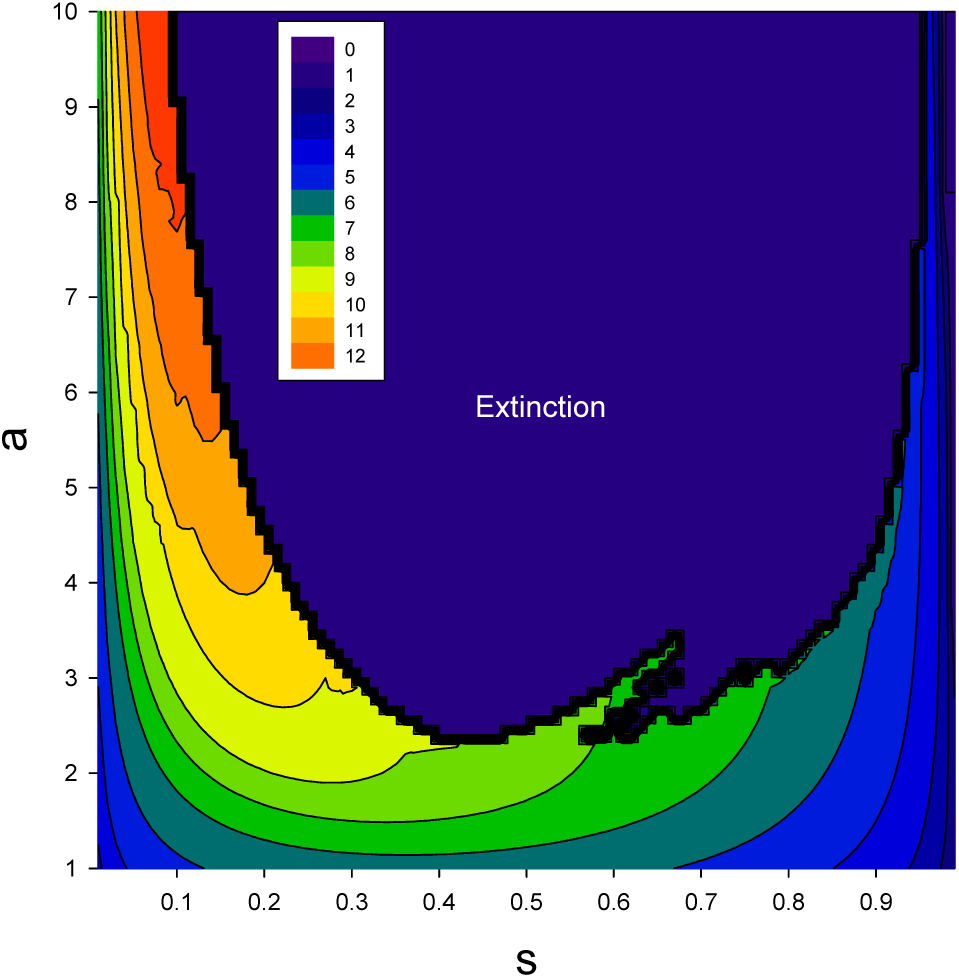
Amplitude profile of stable oscillations through this region of parameter space when the model is synchronized to an oscillation of period 9.9 generations and amplitude of 0.025. Legend indicates *ln*(*amp*) values by color.

**Figure 9.**
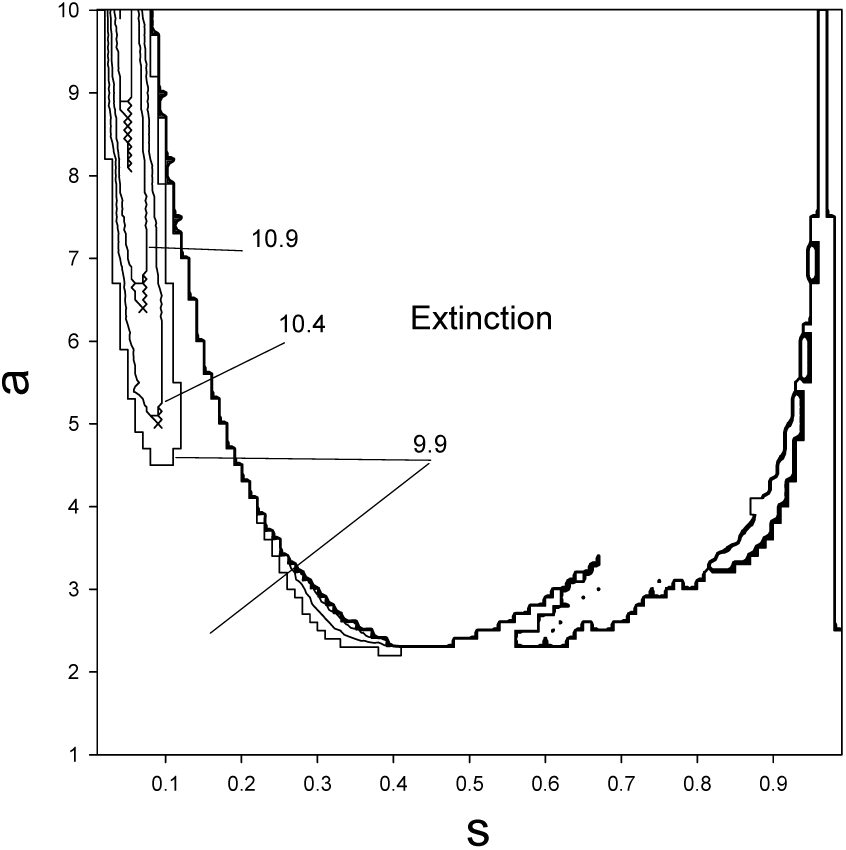
Profile of cycle periods (in generations) within the stable oscillatory region of the parameter space This example was generate with a sinusoidal forcing oscillation of period 9.9 and an amplitude of 0.05. Contour lines are provided at 0.5 intervals. Oscillation periods vary smoothly across the stable region where contour lines exist.

### 5.4 Comparison to wildlife cycles

The frequency distribution of various population cycles in nature is shown in Figure 10. This select set of data was compiled by Anderson & Gillooly (2017) for their study examining cycle periods as a function of the lifeform traits of generation time, body mass and body temperature. Only cycle periods considered ‘significant’ in their Appendix were used here. A 3-point binomial smooth was used on the frequency distribution to reduce fluctuations. Two dominant peaks appear near 6.5 yr and 10 yr. These peaks are dominated by cases that have 1-year generation intervals. These cycle periods are very similar to the maternal and grandmaternal model predictions.

**Figure 10.**
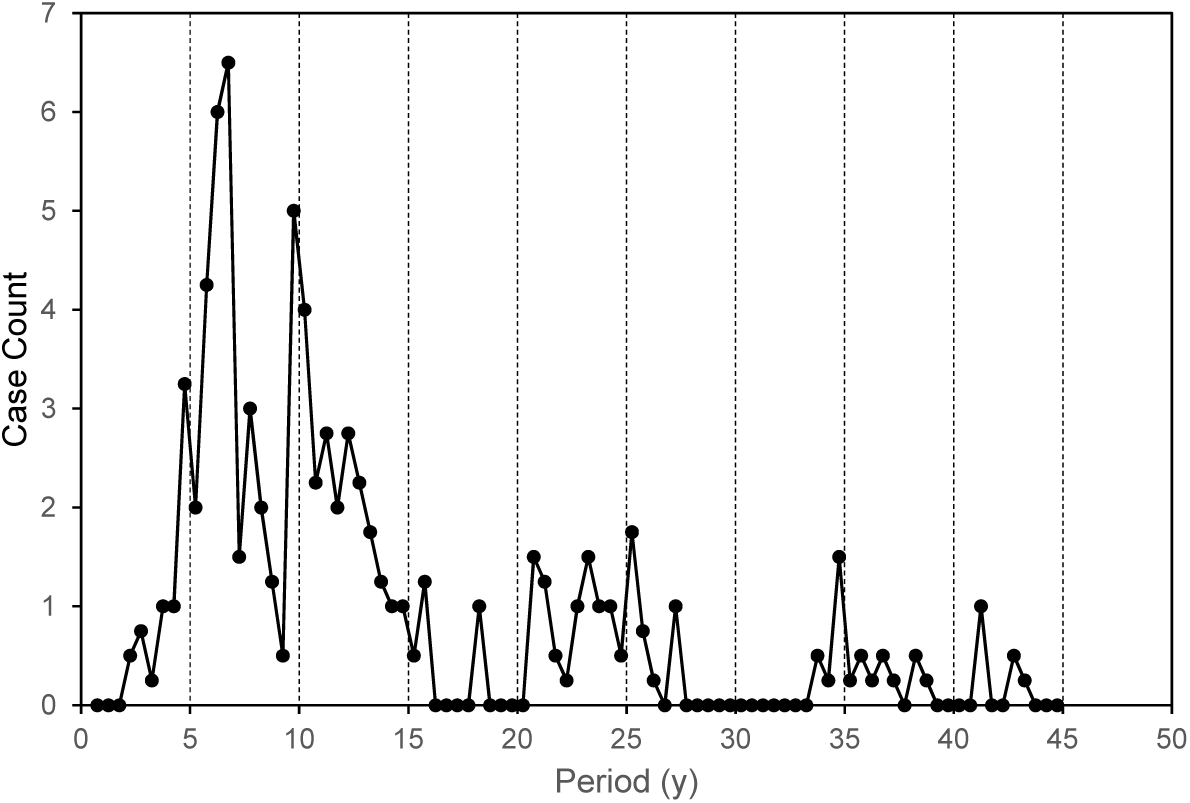
Frequency distribution of wildlife cycle periods compiled by Anderson & Gillooly (2017) and provided in their appendix. These cases included only those deemed statistically significant by the original authors. A 3-point binomial smooth has been applied to the initial frequency distribution to reduce fluctuations.

Focusing on the two-generation delay model and using Eq. (13) it is possible to estimate the cycle length of oscillating populations using only the time between generations. This is shown in Table 2 for some examples of mammals, birds and insects that exhibit population cycles. In all cases, the simple scaling of the inter-generation time by ∼10 yields close estimates of the observed species cycles. The small mammal predator-prey wildlife cycles have been demonstrated many times in the literature and there is little doubt of their cycling. Evidence for the moth cycles extends back several hundred years in some cases and the cycle periods are robust (see references in Table 2). The univoltine, bivoltine, and hemivoltine species robustly exhibit the cycles predicted by this model.

**Table 2:**
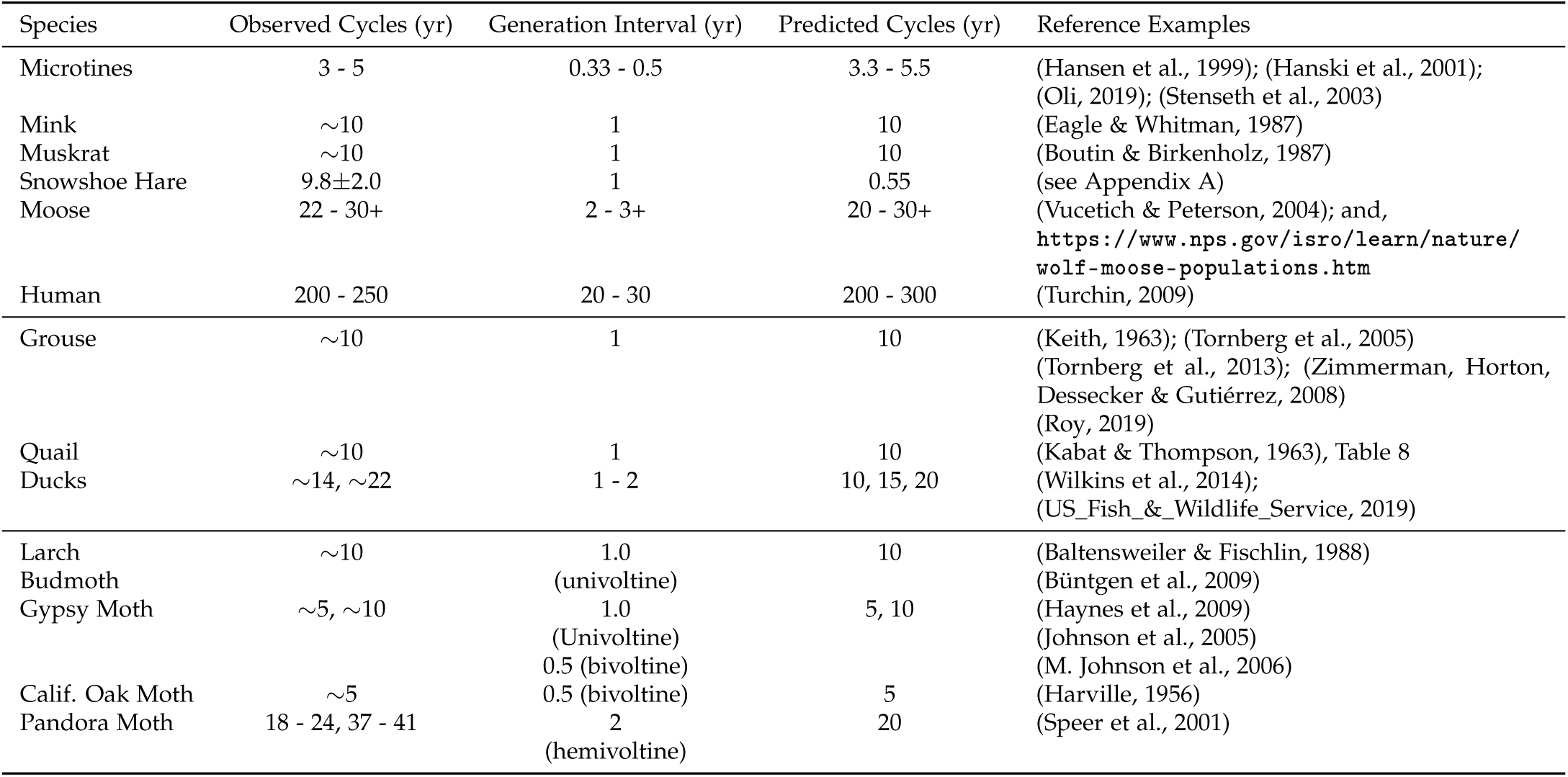
Examples of observed and predicted population cycles.

These examples are also consistent with environmental entrainment by oscillatory components at or near the predicted cycles. An example of possible entrainment is shown in the next Section. Many European moths oscillate with cycles in the 8-10-yr range (e.g., Cerrato et al. (2019)). This could be indicative of very strong environmental signals in this range, such as the North Atlantic Oscillation. It could also indicate a mixture of maternal and grandmaternal epigenetic programming.

### 5.5 Heuristic example of PDO as possible synchronizing signal

Among the examples in Table 2, the snowshoe hare and the ruffed grouse are well-documented, and these cycling populations are located in northern North America. Both have one generation per year and are likely sensitive to the weather and climate of the region (e.g., Cary & Keith (1979); Zimmerman et al. (2008)). One of the drivers of climate in the north and western North America is the pacific sea surface temperatures represented by the PDO time series (Whitfield et al., 2010). This provides an example time series to use as a possible synchronizing function in the grandmaternal model.

The possibility that an oscillating environmental force causes the near spatial synchrony of the hare-lynx cycle throughout Canada has been hypothesized and examined several times (e.g., Krebs et al. (2001); Sinclair et al. (1993)). While this question cannot be answered here, it is shown that this model is consistent with such an effect when driven by an environmental signal related to weather.

The snowshoe hare cycle is described in Appendix A where a mean series is generated from various reports over the years from Alaska and northwestern Canada. The ruffed grouse population series obtained from observations in Minnesota was described in the Methods section. The PDO is shown in Appendix B, Fig. B.1. The Fourier spectral density of the series is shown in Fig. B.2a and the harmonic component that could entrain the model, using the bandpass filter of Fig. 7, is shown in Fig. B.2b.

A portion of the snowshoe hare cycle is shown in the bottom trace of Fig. 11 and the Minnesota grouse cycle is shown in the top trace. The similarity and near synchrony between the hare and grouse cycle is clear, even though the animals in question are separated geographically. The spectral analysis of the snowshoe hare and ruffed grouse time series are shown in Fig. 12a,b and the PDO cycle that could entrain the model, from Fig. B.2b, is shown in Fig. 12c for comparison. The similarity of spectral power is clear, however such similarity should never be construed as cause and effect without much more evidence. These results are provided as a simple example to show that:

**Figure 11.**
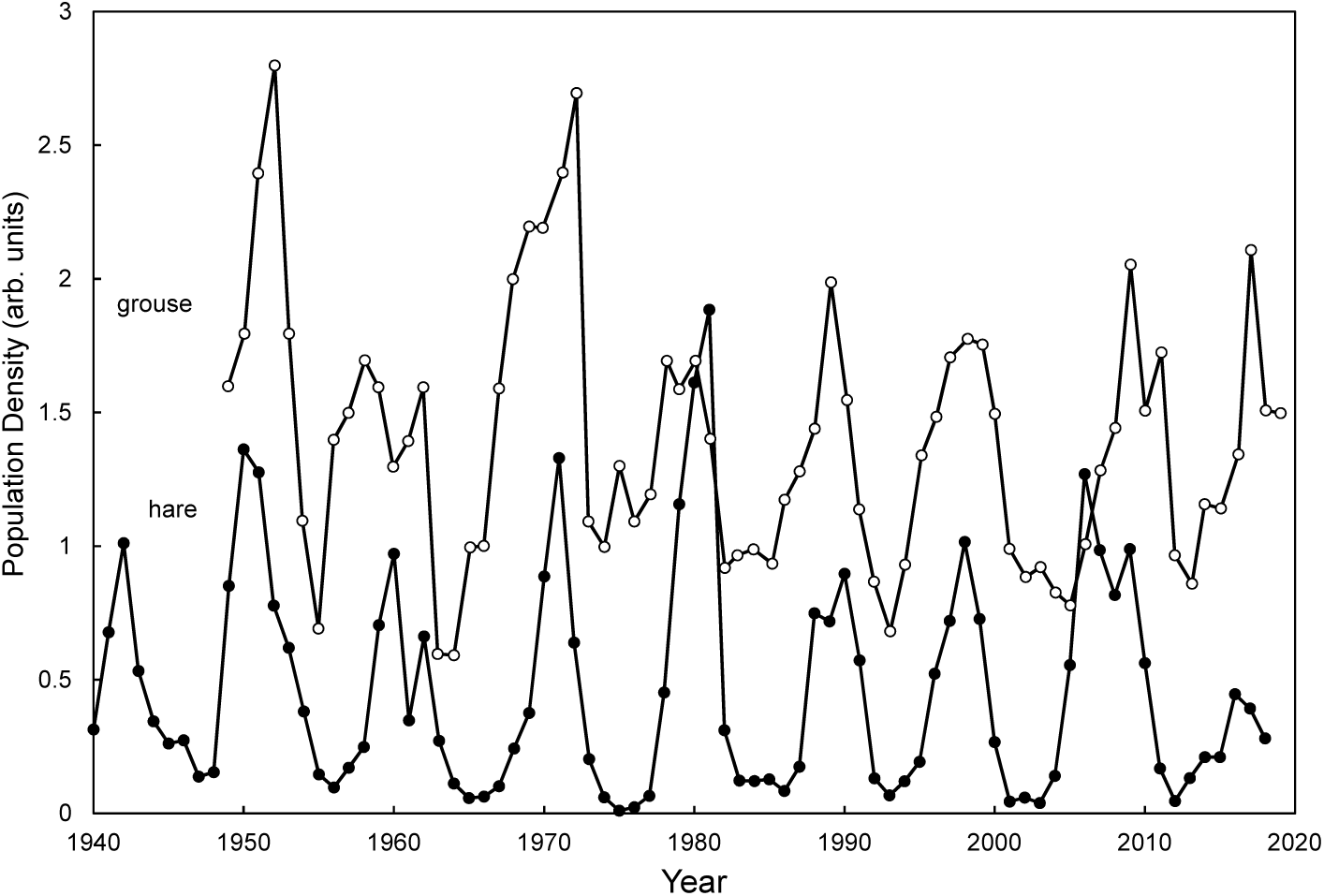
Snowshoe hare composite series (see Appendix A) (solid circles), Minnesota ruffed grouse series (see Methods) (open circles).

**Figure 12.**
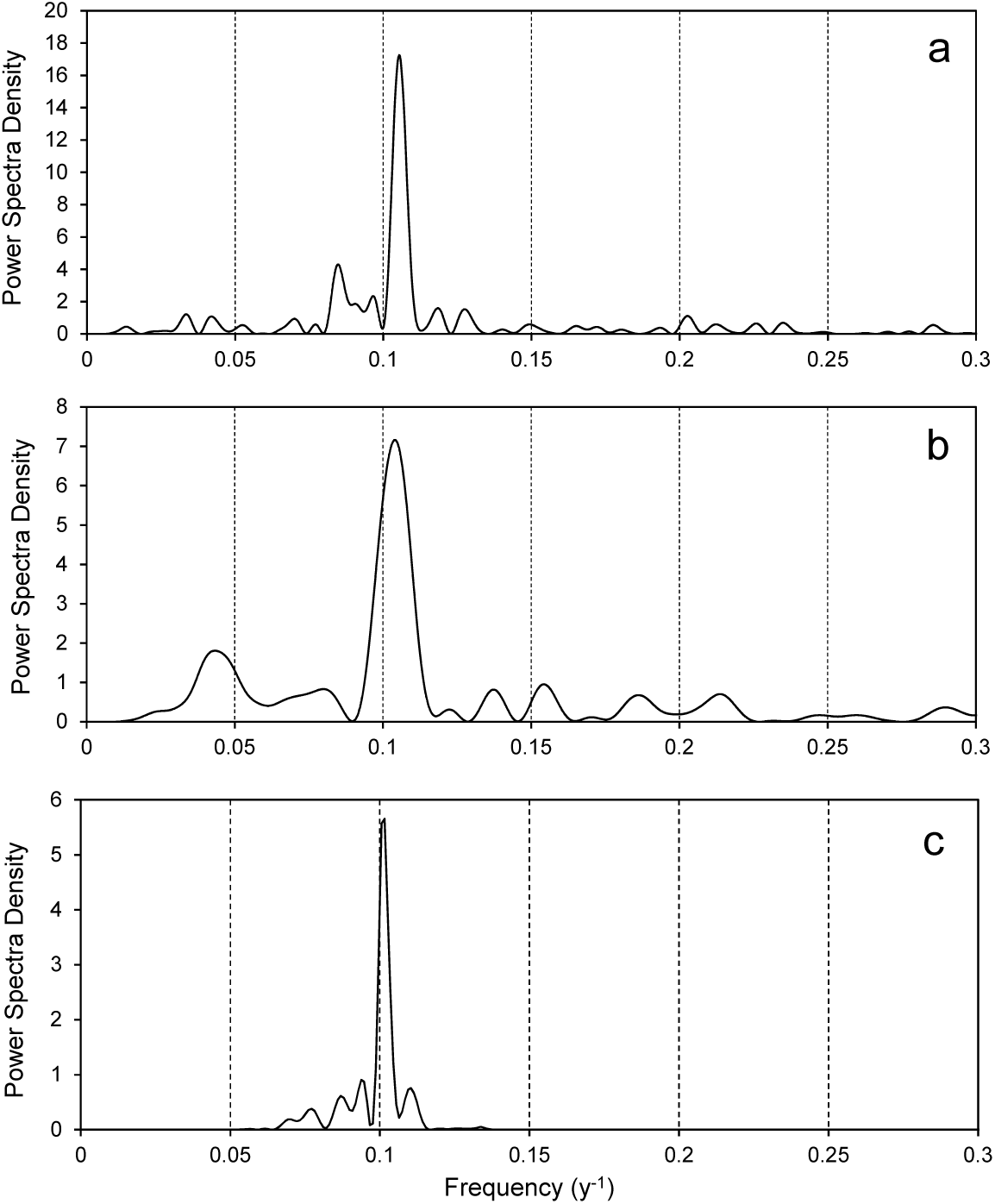
Spectral analysis of the snowshoe hare series (panel **a**), the ruffed grouse series (panel **b**), and the portion of the PDO series that could entrain the model (panel **c**), replotted from Fig. B.2b.

- If wildlife populations are exhibiting the dynamics caused by germline transgenerational signaling, and
- If an environmental signal has the appropriate frequency and amplitude, and
- If the species are sensitive to the environmental signal,
- Then such an signal could synchronize susceptible populations in a manner as shown in Fig. 11.

## 6 Discussion

### 6.1 Overview

First and foremost, this is a hypothesis paper. The five hypotheses outlined in Section 2.2 can be summarized as presenting an approach to harmonize transgenerational epigenetic effects with population dynamics. The postulates and hypotheses have been rendered in a population growth model using a minimum of parameters in a canonical formulation. Both somatic and germline transmission across generations was modeled. Examination of model dynamics reveals spontaneous cycling within regions of the model parameter space for both forms of transmission. The germline transmission (grandmaternal effect) model was examined in more detail and shown that the model can be entrained in both frequency and phase when the external frequency is near the natural oscillation of 10-generations. This version of the model seems consistent with many wildlife cycles across mammals, birds, and insects. As with any hypothesis, testing is required to resolve the degree of validity.

The mathematical model was specifically formulated to contain a minimum number of parameters to allow a comprehensive examination of its behavior as a natural oscillator and as a forced oscillator. A limited set of results were presented to highlight model stability, dynamic range, fundamental frequency range, and reaction to external oscillations.

Many investigators have explored mechanistic models that are tailored to individual species and their niches or to their constellation of prey (e.g, Matthiopoulos et al. (1998); Dobson & Hudson (1992) for the red grouse; Wang et al. (2009)) for predator maturation). These, and many other specialized models, yield cycling dynamics for the species under investigation. The purpose of the model presented here is not to replace or diminish the importance of mechanistic models but to propose that a universal underlying biological mechanism may be at work that predisposes species’ populations to cycle as a consequence of their dynamic epigenetic adaptations to changing environments. Mechanistic models will likely be needed to describe the unique character of each cycling species.

### 6.2 Comments on density-dependent term

While the formulation of Δ in Eq. (5) uses an exponential term, it can be replaced with the simple ratio, such as the one given by:

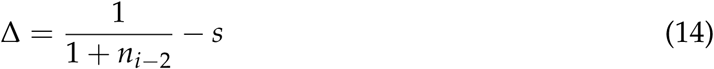

or a logistic equation with the form:

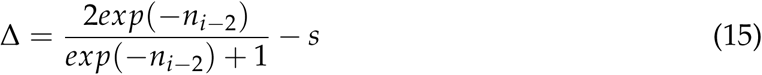

Both the exponential and the ratio terms within Δ provide rapid change at low *n* values and saturation at high *n* values. Regardless of the form these terms take, the introduction of the set point, s, causes the model to operate within the domain below saturation and above zero, thus these limits are never reached in a stably oscillating population. The stability regions within the [*a, s*] parameter space shift somewhat when using the form of Δ in Eqs. (14 or 15), but the period is still robustly near 10 generations. Similar to Eq. (6), the scaling factor *K* can be incorporated in any of the logistic-type formulations to match the model to any given population size.

### 6.3 Comments on forced oscillations

Forcing oscillations are considered to originate externally to affected species. Examples might be weather-related phenomena, primary predators or dependent prey with dominating oscillatory population dynamics, external but nearby predator-prey cycles influencing the local environment, space weather changes in background radiation or magnetic fields, or intermittent pathogen epidemics. All of these typically have complex dynamics that can be deconstructed using harmonic functions as typically done with periodograms and Fourier analysis.

Harmonic deconstruction yields power in various frequency bands whether a series is stationary, near-stationary, or completely chaotic. (See Canyelles-Pericas et al. (2017) for an example in a chaotic system.) Regardless of the nature of the series, these harmonic components can be considered real when the series pumps another oscillatory system with a natural resonance. A resonant system extracts energy from the forcing function in a similar fashion to FFT decomposition. It would do so by extracting instantaneous energy from a component signal. For example, the ∼10-yr component of the PDO yields a constant amplitude harmonic when the whole series is decomposed by FFT, but when the series is examined by wavelet analysis (not shown) the ∼10-yr cycle is strong from 1950 to 2018 and again before 1900. While the phase remains stable throughout the PDO series the ∼10-yr component has varying amplitude. This would be predicted to affect pumped oscillators, such as wildlife cycles, by introducing amplitude modulation.

Noisy environments can also contribute to amplitude modulation (Greenman and Benton, 2005) because they will contain frequencies near the resonant frequency of a natural oscillator. This is a form of stochastic resonance (Wang et al., 2018).

### 6.4 Comments on parameter stability and cycling

Slight shifts in stress or fertility can shift the dynamical model to a region of instability in the [*a, s*] parameter space either temporarily or permanently. This could cause a species to move into a stable, non-cycling region, change its fundamental oscillatory frequency, or become extinct locally. These may be considered tipping points for the species locally (Moore (2018); Pruitt et al. (2018)). The first two of these shifts could be short lived leading to the often observed cycling hiatus occurring during some time periods. Examples of these have documented for the gypsy moth by Allstadt et al. (2013) and in laboratory grown flour beetles by Henson et al. (2003).

### 6.5 Comments on predator dynamics

When the grandmaternal effect leads to cycling in prey species, it seems likely that a bottom-up mechanism would entrain predator species that might naturally oscillate with a different time period. While not shown in this report, the model exhibits harmonics of the native cycling frequency in certain regions of the *a, s* parameter space. As an example, lynx have an inter-generational period of approximately two years, yielding a native cycle of approximately 20 years for values of *a* and *s* in the oscillatory domain. This cycle is overwhelmed, however, by the huge amplitude of the hare cycle creating a ‘bottom-up’ effect operating on the lynx. This is driven by the basic Lotka-Volterra predator-prey cycling (May, 1972). However, the prey also tends to drive the predator cycle at the first harmonic of the predator’s intrinsic cycle. This will naturally create beats and some overall amplitude modulation in both species. The bottom-up effect could also affect multiple predators when a predator hierarchy exists (Flagel et al., 2016).

### 6.6 Comments on time versus generation

Focusing on generation intervals simplifies comparisons among species, which is particularly true when generation intervals are well-defined over long periods of time. For species with intermittent changes in the generation interval, such as cycling rodent populations in the far northern climes with the possibility of two to three generations per breeding season, the population dynamics becomes a complex mixture of possible cycle periods in the time domain. This can cause sequential cycles of varying length and amplitude (not shown) and it is postulated that this will be more common in areas where there are large year-to-year fluctuations in weather variables (e.g., temperature, rain, snowpack, etc.). Therefore, the complexity of the time domain can be simplified conceptually by considering the generation domain even though mechanistic models are still required to explain the variations in generation number in a given season.

This time complexity can readily be seen in the case of many forest defoliators that have both univoltine and bivoltine reproduction (Haynes et al., 2014). With either one generation versus two generations per year, the time domain can yield combined ∼10-yr and ∼5-yr cycles (e.g., Haynes et al. (2009)), which in turn are likely to be entrained to nearby cycles by external forces with similar periods. While working in the generation domain can help simplify the dynamics it cannot account for all variation or for species with no consistent cycling. This is likely due to all the trophic factors that influence population dynamics, as in the case of some forest defoliators (Haynes et al. (2013); (Haynes et al., 2014); Johns et al. (2016)).

### 6.7 Implications

This hypothetical model, if validated, also has implications for species evolution. The set point, *s*, defines an equilibrium that is set by various conditions in the environment and is detected by the reproducing members of the species. It is postulated that the detection of the set point is a genetically determined characteristic of the species, which has evolved to thrive at that equilibrium setting. When the environment permanently changes, resulting in the need to recognize a new set-point, the species must evolve accordingly. This can most readily occur if an epigenetic transgenerational plasticity already exists to provide the mechanisms for population expansion or contraction. This allows natural selection to act rapidly, first to select the epigenetic changes that can become inheritable imprints and then to select genetic mutations that may eventually supplant the imprints.

Following this line of argument, there is an interesting speculation that one can make about cycling. Besides cycling being a natural outcome of the maternal and grandmaternal effects, cycling might allow the population to briefly expand and together with the Allee effect (Allee (1931); Stephens et al. (1999)) provide more fitness in a larger number of individuals to allow for more robust natural selection.

## Abbreviations

PDO: Pacific Decadal Oscillation
NAO: North Atlantic Oscillation
SSTA: Sea Surface Temperature Anomaly
FFT: Fast Fourier Transform
RMS: Root Mean Square

## Acknowledgements

The author is supported by Michigan State University. This research did not receive any specific grant from funding agencies in the public, commercial, or not-for-profit sectors. This work benefited from past communications with Dr. Tim Benton, Dr. Linus Svensson, Dr. Lev Ginzburg and Dr. Eric Kasten.

## A Appendix - Compilation of Snowshoe Hare Series

Data of the snowshoe hare cycles were obtained from several sources. The hare pelt series (1849-1904) was obtained from Maclulich (1957). The hare abundance surveys compiled by Keith (1963) were transformed into a normalized series. Specifically, abundance and scarcity reports were averaged for each year from 1902 to 1948. Abundance values were assigned positive numbers and scarcity values negative numbers. This series was then normalized to fall within the range from zero to one. Counts from local regions were taken from the compilations of Keith (1963), Keith & Windberg (1978) plus field study measurements of Cary & Keith (1979), Keith (1990), Krebs et al. (1995), Alaskan data from wildlife biologist Steve DuBois (Tim Mowry, “Snowshoe Hare Peaks in Interior Alaska”, Alaska Fish and Wildlife News, Alaska Department of Fish and Game, Dec. 2007, http://www.adfg.alaska.gov/index.cfm?adfg=wildlifenews.view_article&articles_id=339), Merizon & Carroll (2019), and Krebs et al. (2018). A compilation series was constructed by scaling each separate series by its root mean square (RMS) amplitude, which allowed comparison across various types of observational units. While such scaling precludes extracting meaningful amplitude variations, it allows a suitable series for detecting the primary frequency component. The various series and the composite series are shown in Fig. A.1.

**Figure A.1.**
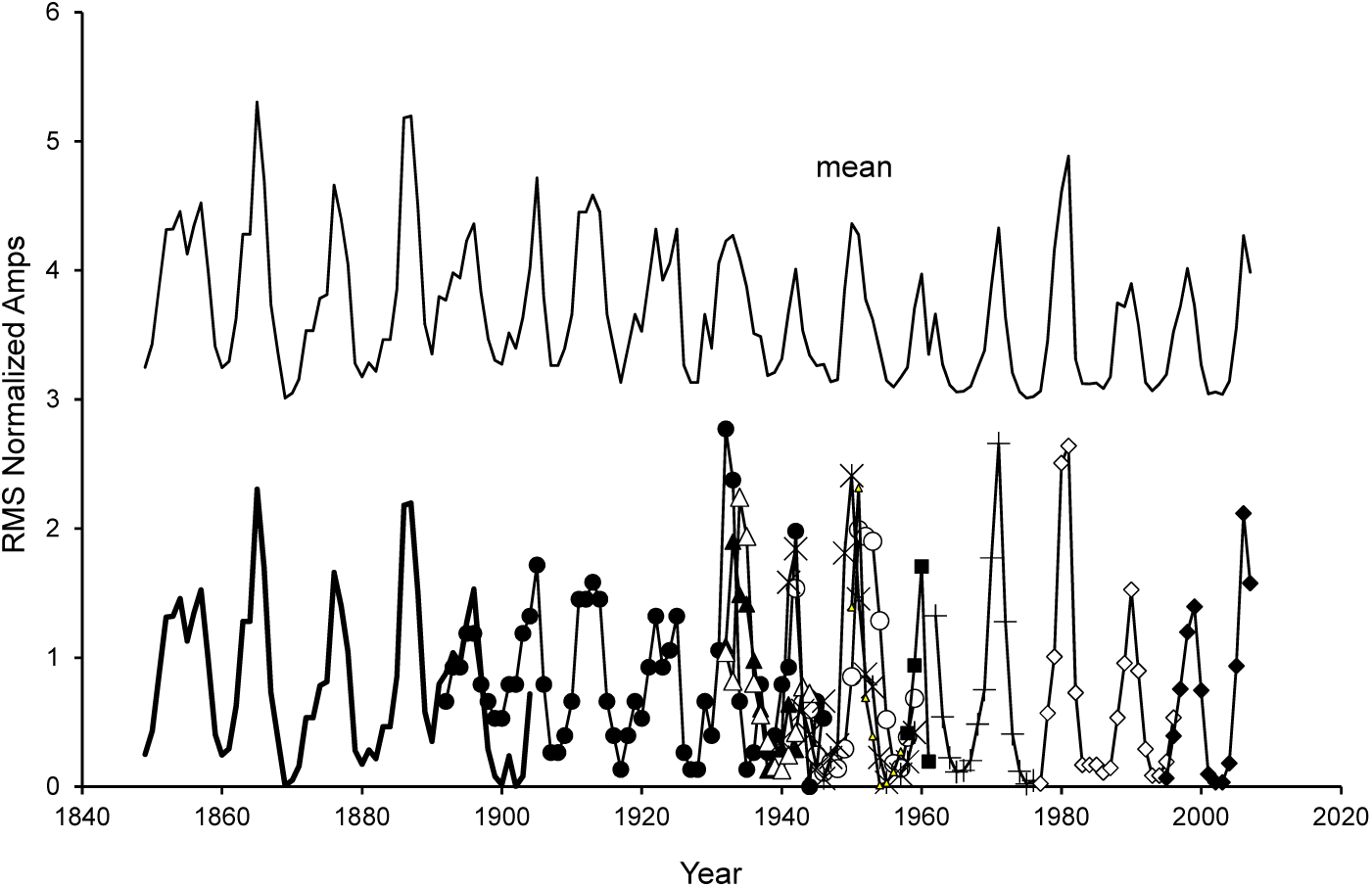
Compilation of snowshoe hare population observations from various sources. (Lower Panel): RMS normalized time series from various sources: (solid line) Hudson bay series, (Maclulich, 1957); (solid circle) Hare abundance surveys (Keith, 1963); Small area counts, (open circle) Ontario, (X symbol) Minnesota, (open diamond) Wisconsin (Keith, 1963); (+ symbol) Alberta (Cary & Keith, 1979); (open triangle) Yukon (Keith, 1990); (bold dashed line) Yukon (Krebs et al., 2018); (solid triangle) Alaska (Wildlife News Alaska, 2007); (open squares) Denali Alaska (Merizon & Carroll, 2019), (asterisks) Delta BBS Alaska (Merizon & Carroll, 2019). (Upper Panel) Average of RMS normalized series of the lower panel plus a few more short series from (Keith, 1963) and from (Keith & Windberg, 1978). The upper panel is offset vertically for clarity.

## B Appendix - Pacific Decadal Oscillation & 10-yr Component

The global sea surface temperature anomaly (SSTA) series has been recorded for several decades. A component of that series is the Pacific Decadal Oscillation (PDO) (see Methods). The detrended version of this series is shown in Fig. B.1. The Lomb Periodogram spectral analysis of this is shown in Fig. B.2a. The portion of that spectrum that is capable of driving the model utilizing Eq. (12) is shown in Fig. B.2b. This was obtained by using the bandpass amplitude profile of Fig. 7b as an envelope to extract the PDO band most likely to entrain the model.

**Figure B.1.**
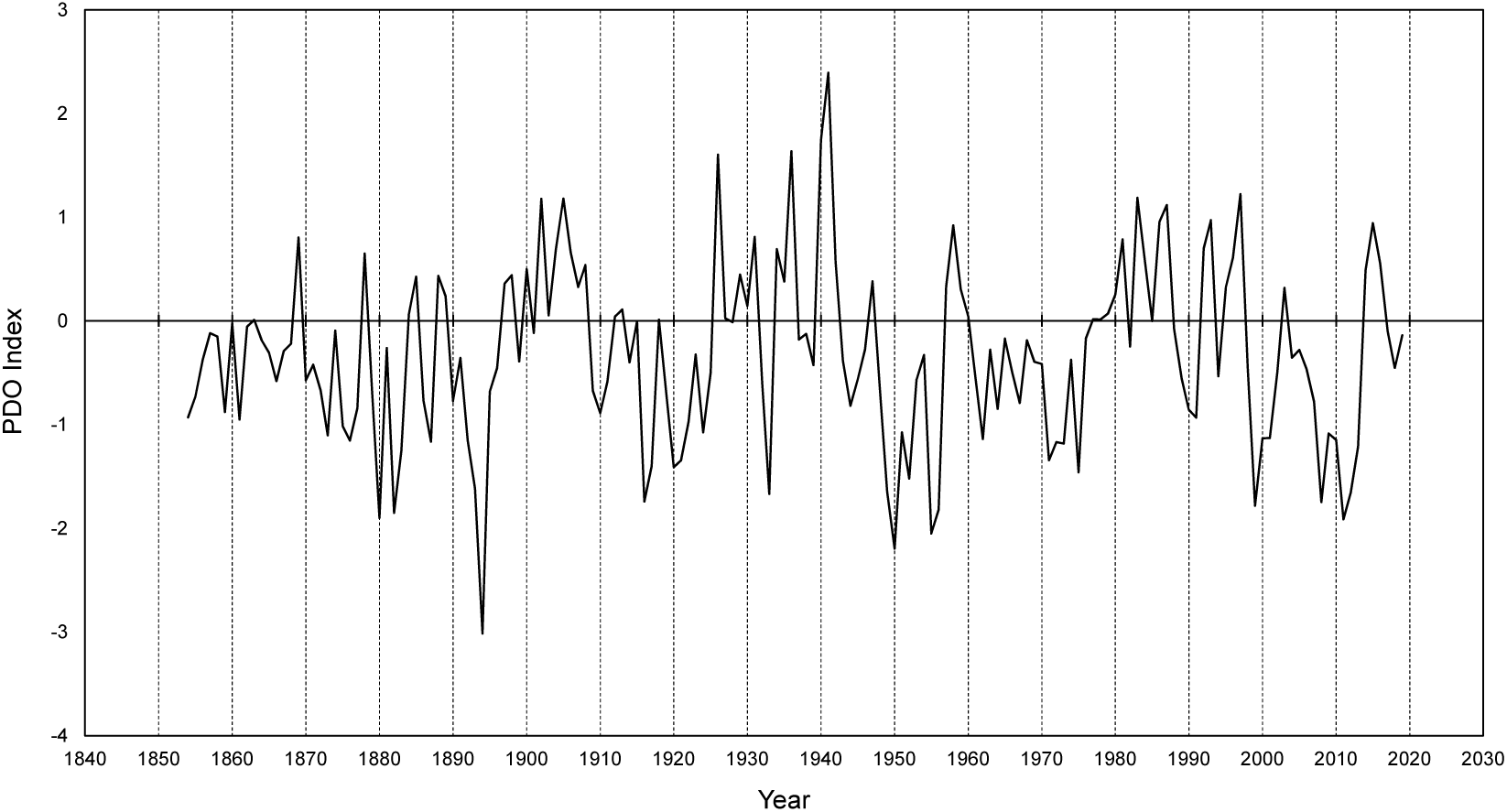
Pacific Decadal Oscillation index (see Methods) detrended with a 2nd order polynomial. The PDO was obtained as a monthly series and was converted to yearly spaced values by averaging over each calendar year.

**Figure B.2.**
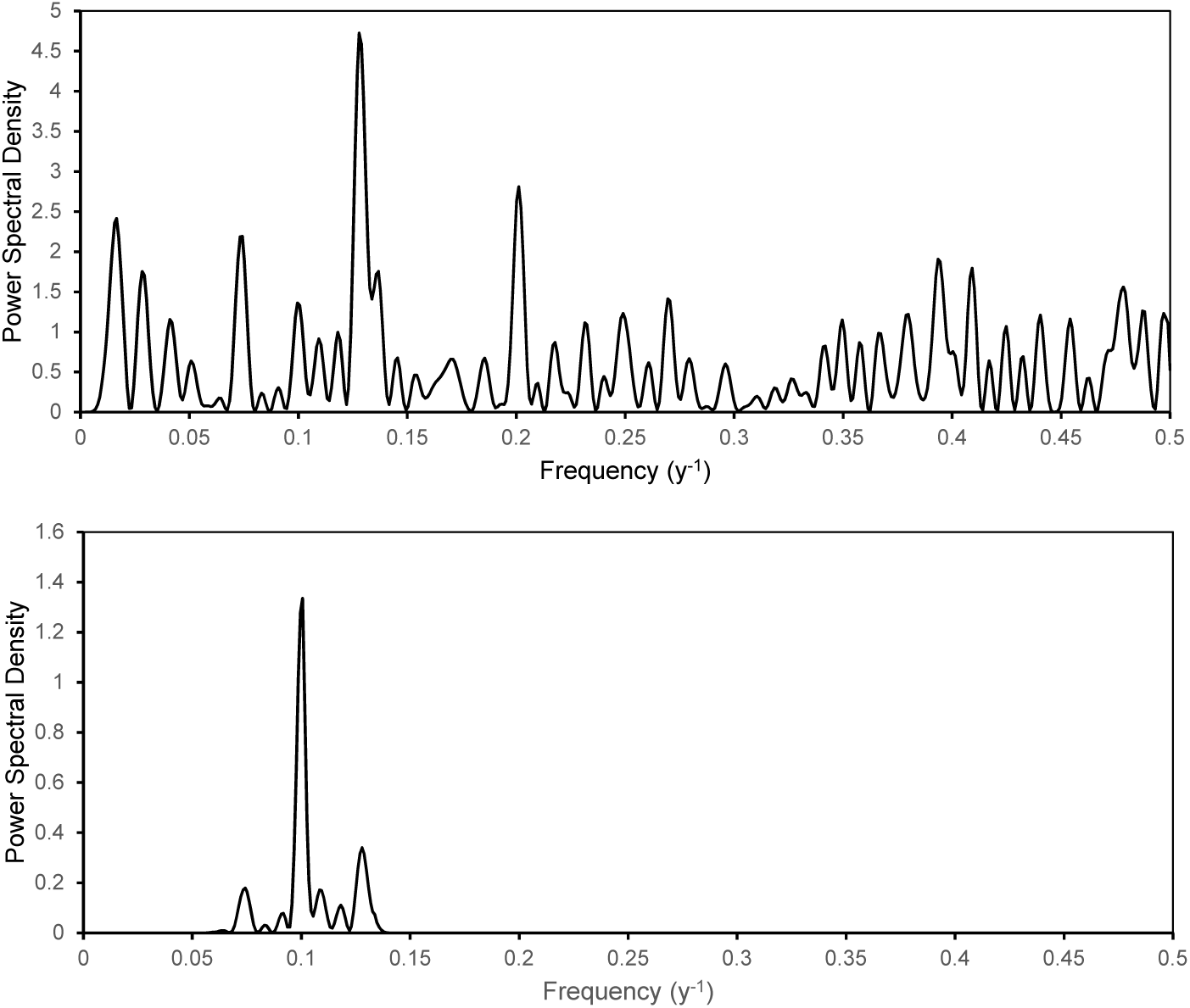
Lomb periodogram spectra of the PDO. (*top*) full spectrum of the PDO. (*bottom*) full spectrum multiplied by the ‘bandpass’ envelope of Fig. 7b. Regions beyond the envelope are given values rapidly approaching zero.

